# Does Neuroticism Cause Depression? A Mendelian Randomization Study

**DOI:** 10.1101/420703

**Authors:** Doug Speed, Gibran Hemani, Maria S. Speed, Major Depressive Disorder Working Group of the Psychiatric Genomics Consortium, Anders D. Børglum, Søren D. Østergaard

## Abstract

Neuroticism is a heritable personality trait, characterized by negative emotions such as worrying, feelings of guilt, loneliness and being easily hurt. Increased levels of neuroticism are associated with poor mental health – development of depression in particular – but it remains uncertain whether this association represents a causal effect [1]. Here, we use Mendelian randomization (MR) to investigate whether neuroticism is a causal risk factor for development of depression.

MR is an analytic approach to assess the causality of an observed association between a risk factor and a clinically relevant outcome [2]. MR is particularly useful in situations where randomized controlled trials are not possible and observational studies provide biased associations because of confounding or reverse causality. MR uses genome-wide association study (GWAS) data to identify instrumental variables (single nucleotide polymorphisms; SNPs) for a risk factor (here neuroticism) that are then tested for association with the outcome of interest (here depression). MR exploits the fact that SNP genotypes are randomly allocated during gamete formation (Mendel’s second law) and are thus generally not susceptible to reverse causation and confounding [2]. Therefore, MR is often referred to as “nature’s randomized control trial”.

For this MR study we used summary statistics from the largest published GWAS of neuroticism (449,484 samples) where individuals were scored based on the 12 neuroticism items from the Eysenck Personality Questionnaire (EPQ) [3], and from the largest published GWAS of major depression (135,458 cases and 344,901 controls), where diagnosis was based on either self-reporting, clinical assessment, or examination of medical records [4]. We identified 82 independent, genome-wide significant SNPs for neuroticism; starting with the 7,759 genome-wide significant (P<5e-8) SNPs from the neuroticism GWAS, we excluded ambiguous SNPs (those with alleles A&T or C&G) and those not present in the depression GWAS, then thinned so that no pair of SNPs within 3 centiMorgan had correlation-squared >0.001. Collectively, the 82 SNPs explain 0.89% of the variation in the EPQ neuroticism score.

Figure 1 shows that for the 82 SNPs, there is a positive correlation between effect sizes for neuroticism and depression. Using inverse-variance weighted regression (red line), as implemented in the R package MR-Base [2], we estimated the slope to be 0.25 (SD 0.02), which is significantly positive (P<1e-16), and therefore strong evidence that neuroticism is a causal risk factor for depression. Specifically, these results indicate that every additional YES answer to the neuroticism items in the EPQ corresponds to a 0.25 higher log odds ratio (OR) for depression (i.e., a 1.28-times higher OR).

**Figure 1.**
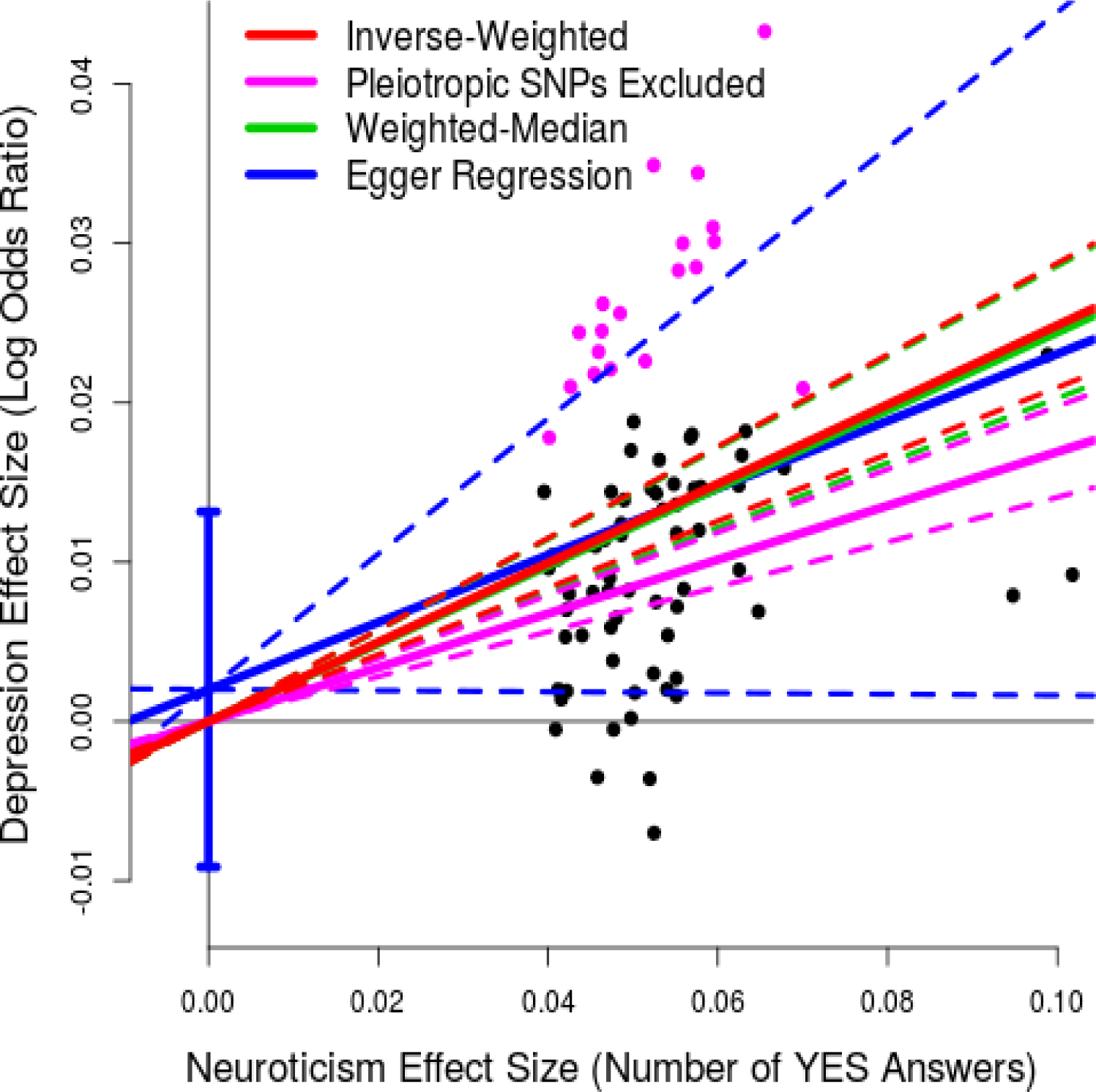
Results of Mendelian Randomization (neuroticism to depression) For 82 independent, genome-wide significant (P<5e-8) SNPs for neuroticism, points report per-allele effect sizes for neuroticism and depression (the units are number of YES answers to the 12 neuroticism items on the Eysenck Personality Questionnaire and log odds ratio, respectively). The solid red line indicates the estimated slope from inverse-variance regression (the solid purple line indicates the slope if the purple SNPs, those nominally associated with depression, are excluded). The solid green line indicates the estimated slope from weighted-median regression; the solid blue line indicates the estimated slope from Egger Regression (the vertical blue segment marks a 95% confidence interval for the intercept). The four pairs of dashed lines mark 95% confidence intervals for the slopes.

Underlying MR are three key assumptions [2], which in the context of this study are: (i) all 82 SNPs are associated with neuroticism; (ii) the 82 SNPs are uncorrelated with confounders of the neuroticism-depression association; (iii) the 82 SNPs affect depression only through neuroticism and not directly (no pleiotropy). Our decision to use only SNPs robustly-associated (rather than putatively-associated) with neuroticism should ensure (i) is true, while (ii) should be satisfied because each individual’s genotypes for the 82 SNPs are randomly assorted during gamete formation. To explicitly test (iii), we would have to perform a conditional analysis for depression (i.e., confirm that each of the 82 SNPs is not associated across individuals who score 0 out of 12 for neuroticism, nor across individuals who score 1 out of 12, etc.). However, this is not feasible, so we instead performed three sensitivity analyses [2], also reported in Figure 1, which confirmed that: the slope remains significantly positive (0.17, SD 0.02) if we exclude the 19 SNPs (those marked in purple) with P<0.05/82 for depression (i.e., those showing strongest evidence for pleiotropy); likewise the slope remains significantly positive (0.24, SD 0.02) if we instead use weighted-median regression (green line), which is robust provided at least 50% of the information comes from non-pleiotropic SNPs; the intercept from Egger Regression (blue line) is consistent with zero (0.002, SD 0.006), indicating there is no strong evidence for directional pleiotropy.

In the supplementary material (Supplementary Figure 1), we provide results from two additional analyses. Firstly, noting that some samples were used in both the neuroticism and MDD GWAS, we verified that the MR results were similar if we repeated the analyses using summary statistics from independent sub-GWAS. Secondly, we reversed the direction of the analysis, to test whether susceptibility to depression is a causal risk factor for neuroticism. Using the same approach as for neuroticism (above), we identified 27 independent, genome-wide significant SNPs for depression, which in total explains 0.35% of susceptibility (measured on the liability scale, assuming a prevalence of 0.14 [4]). Using inverse-weighted regression, we estimated the slope to be 0.90 (SD 0.11; P<1e-16); once more, the slope remained significantly positive if we excluded the 17 SNPs showing evidence for association with neuroticism or if we instead used weighted-median regression, while the intercept from Egger Regression was consistent with zero. This indicates that there is also a causal effect going from depression to neuroticism.

The most clinically relevant result of this study is that MR has provided strong evidence to support our hypothesis that neuroticism is a causal risk factor for depression. The implication of this finding is that reducing the degree of neuroticism will tend to reduce the risk of depression. This is in agreement with the results by Quilt et al. suggesting that reduction of neuroticism is the mechanism by which selective serotonin reuptake inhibitors exert their antidepressant effect [5].

In terms of the mechanism underlying the causal relationship between neuroticism and increased risk of depression, the syndrome defined by the 12 neuroticism items in the EPQ may provide the answer. This syndrome describes a phenotype characterized by high vulnerability to external environmental stressors and/or adverse life events. Whether high vulnerability to stress is indeed driving the causal effect of neuroticism upon the risk of depression could ideally be investigated by gene-environment interaction studies based on longitudinal datasets containing individual-level information on genotype, exposure to external stressors and development of depression.

## Acknowledgments

We thank the research participants and employees of 23andMe, Inc. for their contribution to this study. D.S. is supported by the European Unions Horizon 2020 Research and Innovation Programme under the Marie Skłodowska-Curie grant agreement number 754513, by Aarhus University Research Foundation (AUFF) and the Independent Research Fund Denmark (7025-00094B). G.H. is supported by the Wellcome Trust (208806/Z/17/Z). A.D.B. is supported by grants from The Lundbeck Foundation (R102-A9118 and R155-2014-1724). Data handling and analysis on the GenomeDK HPC facility was supported by NIMH (1U01MH109514-01 to Michael O’Donovan and A.D.B.). High-performance computer capacity for handling and statistical analysis of iPSYCH data on the GenomeDK HPC facility was provided by the Centre for Integrative Sequencing, iSEQ, and Center for Genomics and Personalized Medicine, Aarhus, Denmark (grants to A.D.B.). The PGC has received major funding from the US National Institute of Mental Health and the US National Institute of Drug Abuse (U01 MH109528 and U01 MH1095320). S.D.Ø. is supported by AUFF (AUFF-E-2015-FLS-7-2 and a Jens Christian Skou Junior Fellowship), the Riisfort Foundation, the Lundbeck Foundation (R165-2013-15320) and the Independent Research Fund Denmark (7016-00048).

